# Modeling Withdrawal States in Opioid-Dependent Mice with Machine Learning

**DOI:** 10.1101/2025.07.29.667254

**Authors:** Savanna A. Cohen, Lisa M. Wooldridge, Justin J. James, Jacqueline W. K. Wu, Gregory F. Corder

## Abstract

Understanding opioid withdrawal behaviors in preclinical models is critical to improving therapeutic approaches for opioid use disorder (OUD). However, quantifying these withdrawal behaviors remains a difficult process for researchers, given the subtlety of behaviors and variation across individuals. To overcome these difficulties, we developed a scalable behavioral analysis pipeline using LUPE (Light aUtomated Pain Evaluator), an open-source framework integrating video acquisition, pose estimation, supervised and unsupervised classification, and expert-guided behavior discovery. Mice undergoing naloxone-precipitated opioid withdrawal were recorded and analyzed using DeepLabCut for markerless pose estimation. We hand-annotated withdrawal-specific behaviors, including jumping, genital licking, grooming, and paw tremors, and normal behaviors, including walking, rearing, and being still, using Behavioral Observation Research Interactive Software (BORIS) to generate frame-by-frame ethograms. The annotations and pose data were then imported into Active learning Segmentation of Open field in DeepLabCut (A-SOiD), an active learning platform for behavior classification. A-SOiD successfully detected some behaviors (e.g., grooming and rearing) which were of a longer duration, though other rapid behaviors (e.g., jumping and paw tremors) were inconsistently captured. While no novel behavioral motifs have been discovered yet, ongoing work aims to refine model performance. This LUPE-based pipeline sets the groundwork for standardized, high-resolution behavior quantification and is being applied to additional datasets to investigate whether new components of the withdrawal phenotype emerge across experimental conditions.

## Introduction

Opioid use disorder (OUD) remains one of the most persistent and devastating public health crises in the United States, with over 100,000 opioid-related overdose deaths reported annually (Cho et al., 2020). It is reported that nearly 8.6 million people have misused prescription opioids, excluding those who use non-prescription opioids like heroin (Cho et al., 2020). One of the most difficult barriers to overcome OUD is withdrawal, a syndrome of highly aversive symptoms that emerge following cessation of opioid intake (Wooldridge et al., 2025). These include both physical symptoms (including muscle aches, nausea, and sweating) and psychological symptoms (including anxiety, irritability, and cravings), which are often so severe that they drive continued drug use to avoid discomfort, creating a powerful negative reinforcement cycle that traps a victim in continued use (Wooldridge et al., 2025). In mouse models, opioid withdrawal manifests through a variety of species-specific behaviors, many of which serve as important proxies for withdrawal severity (Gipson et al., 2021). These include jumping (**Fig. 1B**), grooming bouts, paw tremors (**Fig. 1A**), and genital licking (Uddin et al., 2021). Jumping (**Fig. 1B**) is an escape tactic often associated with hyperactivity in stress-related circuits in the brain (Siegel et al., 1975). On the other hand, paw tremors (**Fig. 1A**) reflect somatic nervous system instability (Kim & Im, 2024). It can be reasoned that these behaviors stem from distinct neural circuits. In addition to this, these behaviors vary widely in duration and frequency, which makes them difficult to detect and quantify using traditional video scoring Historically, researchers have relied on frame-by-frame manual scoring of videos to quantify withdrawal behaviors (Bumgarner et al., 2022). However, this method is time-intensive, subject to human bias, and difficult to reliably reproduce across laboratory settings. In particular, scoring a single 30-minute video can take 2–3 hours per observer and is subject to inter-rater variability. Our lab has previously hand-scored hundreds of withdrawal behaviors as part of broader studies on the neural circuitry underlying fentanyl withdrawal (Wooldridge et al., 2025). These efforts required over 150 hours of annotation and approximately $2,500 in labor costs. The ability to automate behavior detection would drastically reduce annotation time and workload. Furthermore, withdrawal behaviors are prone to being misclassified or omitted due to their brief duration. Overall, these barriers have held back the standardization of withdrawal models and therapeutic treatments as a result (Bumgarner et al., 2022). In this thesis, we aim to accelerate the process and increase precision by leveraging machine learning.

**Figure 1.**
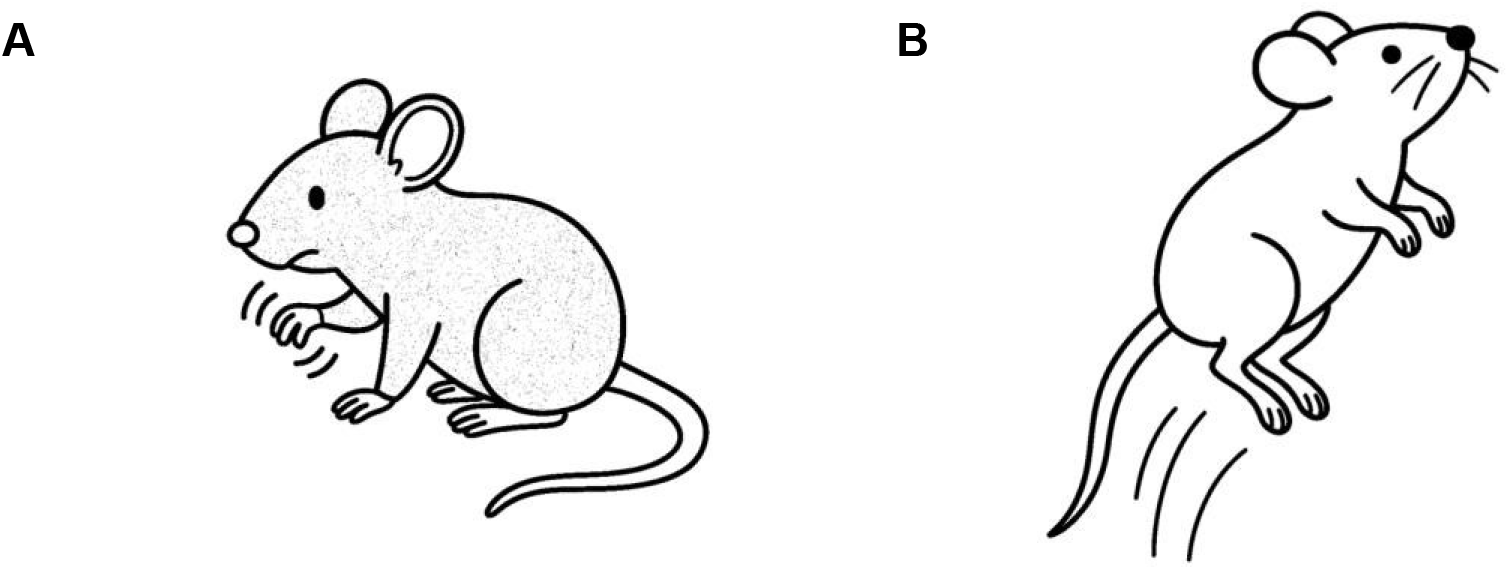
Withdrawal Behaviors in Mouse Models. Mouse models undergoing opioid withdrawal exhibit a range of somatic and affective behaviors. Paw tremors (**1A**) and (**1B**) jumping are hallmark withdrawal signs. Other observed behaviors include grooming and genital licking, which may reflect discomfort or stress regulation strategies. These behaviors form the core behavioral repertoire assessed across subsequent analyses.

We developed a behavioral analysis pipeline that utilizes LUPE chambers developed by our lab. These chambers are standardized open-field arenas that are used to record high-resolution bottom-up videos of various mouse behaviors (Oswell et al., 2024). Bottom-up videos are advantageous as they improve visibility of limb and ventral body movements and minimize visual occlusion (**Figure 2**). The chambers further remove the presence of a human experimenter, allowing the mice to behave in a more natural sense during each experiment.

**Figure 2.**
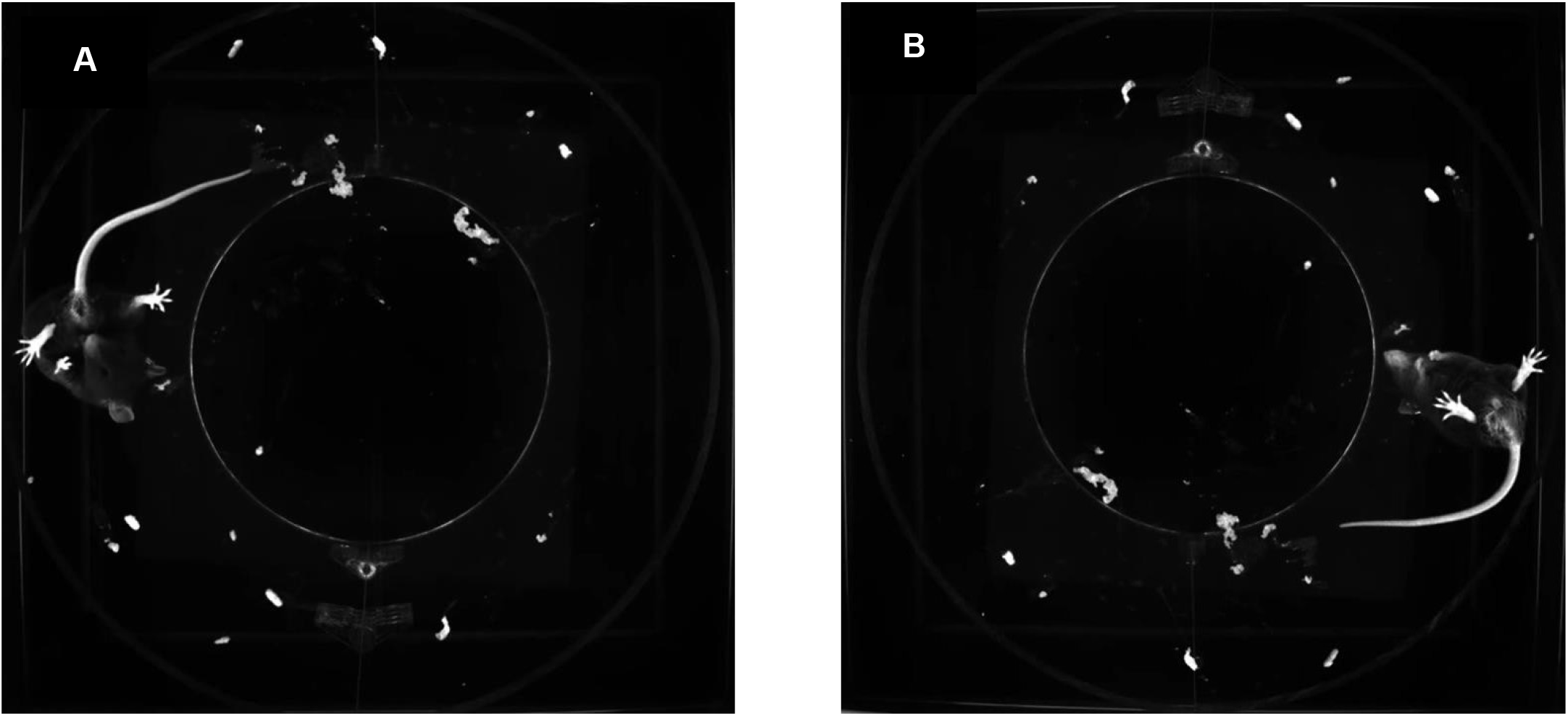
Mouse Behavior Recorded in LUPE Chambers. Representative frames from bottom-up video recordings within the LUPE chambers during a 30-minute post-naloxone observation window in opioid-dependent mice. (**2A**) A mouse displays genital licking—one of the most frequently observed withdrawal-associated behaviors. (**2B**) Rearing, a common exploratory behavior in mice, is also observed and tracked, demonstrating the behavioral diversity captured within this setup.

After videos were collected, we used BORIS, an open-source behavioral annotation tool, to identify and record behaviors and overcome manual scoring limitations (Friard & Gamba, 2016). We further incorporated DeepLabCut, a deep learning-based markerless pose estimation framework, to precisely track movement and behaviors (Mathis et al., 2018). We then tested both B-SOiD and A-SOiD—two open-source machine learning tools developed for behavior classification. B-SOiD, which uses unsupervised learning to segment behaviors based on spatiotemporal patterns in pose data, was our initial approach (Hsu & Yttri, 2021). However, B-SOiD was ultimately excluded from our final pipeline because it was outdated and no longer compatible with our lab’s computing environment, reflecting the broader challenge of maintaining open-source tools that have not been actively supported in recent years.

Nevertheless, we input the data we obtained from BORIS and DeepLabCut into A-SOiD, a machine learning program that combines supervised learning and unsupervised learning with an active learning tool, allowing expert users to iteratively correct uncertain predictions (Tillmann et al., 2024). A-SOiD functions by asking the expert to annotate and refine only examples that it is uncertain about, significantly reducing the burden of manual annotation (Tillmann et al., 2024). Although supervised methods are efficient in identifying known behaviors, using them in conjunction with unsupervised methods can uncover subtle patterns that human observers may miss. When these are integrated with active learning, this strategy allows models to iteratively refine both familiar and novel behaviors with expert feedback (Tillmann et al., 2024).

Overall, we implemented an integrated LUPE–BORIS–DeepLabCut–A–SOiD pipeline to study and record opioid withdrawal behaviors in mouse models. We hypothesized that this pipeline would allow for accurate and reproducible classification of known and novel withdrawal behaviors in mice dependent on either morphine or fentanyl.

## Methods

### Subjects and Drug Administration

All drug administration procedures were completed prior to the start of this project; we received and analyzed video recordings of withdrawal behavior collected immediately following naloxone administration in both morphine-and fentanyl-treated mice.

Adult male and female mice were housed on a 12-hour light/dark cycle with ad libitum access to food and water. To induce morphine dependence, animals received morphine via their drinking water for 14 days with escalating concentrations: 0.3 mg/mL from days 1–3, 0.5 mg/mL from days 4–5, and 0.75 mg/mL from days 6–14. On the final drinking day, naloxone (1 mg/kg) was administered through an intraperitoneal injection to precipitate withdrawal.

In a separate cohort, fentanyl dependence was induced by replacing drinking water with 0.02 mg/mL fentanyl citrate solution for 8 consecutive days. On the final drinking day, naloxone (1 mg/kg) was administered through an intraperitoneal injection and precipitated withdrawal behaviors were recorded (Wooldridge et al., 2025).

It is important to note that morphine and fentanyl exposure paradigms differed in duration of administration, resulting in distinct withdrawal timepoints and behavioral profiles. Both cohorts of mice were recorded for 30 minutes.

### LUPE Behavior Recording

Mice were recorded in LUPE open-field chambers (254mm x 305mm) (**Fig. 3**). Monochrome infrared sensors were fitted 300mm below the glass floor arena to capture 60 frames per second of behavior (Oswell et al., 2025). LEDs were set to 580nm infrared light to create a standardized dark, safe environment for the mice to minimize disturbances to their nocturnal behavior (**Fig. 3**) (Oswell et al., 2025). The bottom-up configuration allowed for consistent capture of limb movement, ventral features, and tail motion, which are key for identifying subtle behaviors like paw tremors and genital licking. The LUPE setup was chosen over top-down systems due to its superior resolution of fine motor movements and better compatibility with pose tracking tools like DeepLabCut.

**Figure 3.**
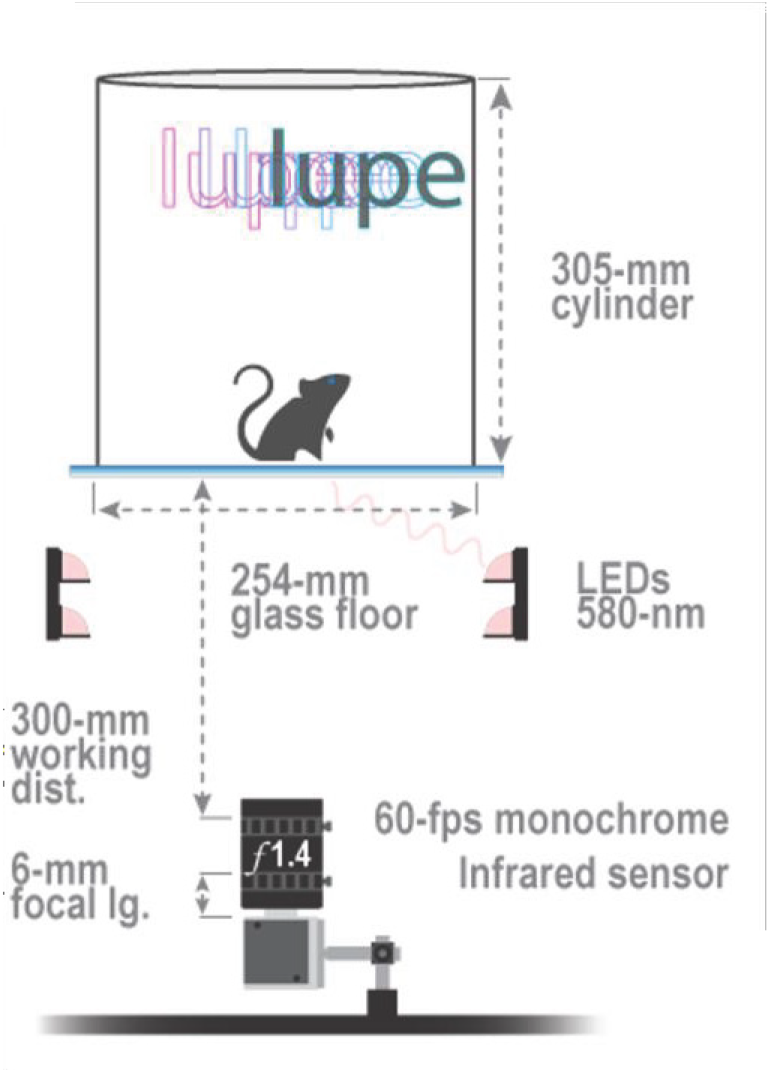
LUPE Behavioral Imaging Setup. Schematic of the LUPE recording environment. Mouse behavior is captured using a 60 frames-per-second camera mounted 300 mm beneath a transparent floor. Side-mounted 580-nm LEDs provide consistent illumination. Infrared sensors eliminate visible light stressors while enabling precise pose estimation through bottom-up imaging (Oswell et al., 2024).

### DeepLabCut: Pose Estimation

We used our lab’s pre-trained DeepLabCut model to analyze recordings (**Fig. 4A**). The network used a 20-body point system to extract 2D coordinates of multiple body parts along with a confidence score (**Fig. 4A**) (Oswell et al., 2024). The data collected included the position and movement of body parts over time which can be used to calculate velocity, inter-limb distances and angles, movement frequency and relative positions in the arena (**Fig. 4B**) (Mathis et al., 2018). The model output pose data and time-aligned kinematics which form the input for downstream behavior classification. This transformation enabled A-SOiD to interpret motion patterns independent of animal size or absolute arena location.

**Figure 4.**
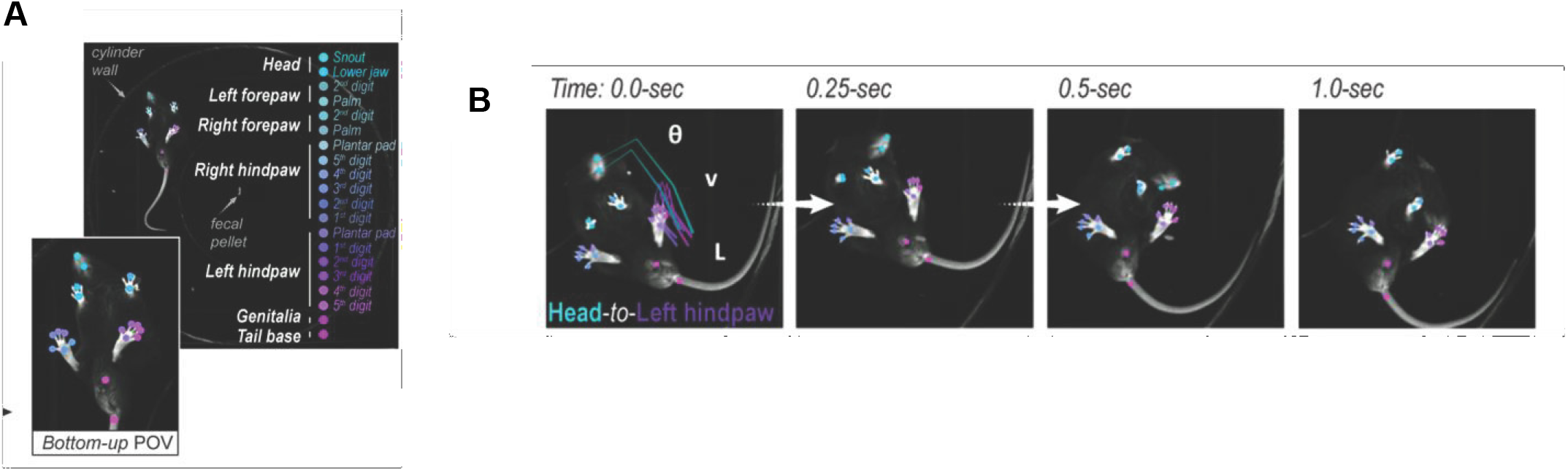
Pre-trained 20 Body-Point Network. DeepLabCut-enabled pose estimation using a pre-trained 20-point body tracking network allows for detailed analysis of movement and posture (**4A**). Keypoints include the limbs, spine, and tail base (**4B**). Time-lapse sequences reveal postural transitions, such as changes in the angle between the head and left hind paw, used to infer specific behaviors (Oswell et al., 2024).

### BORIS: Manual Annotation

Withdrawal behaviors were manually scored using BORIS to develop a dataset for reference standards. The data included ethograms of natural mouse behaviors (walking, being still, and rearing) and withdrawal behaviors (jumping, genital licking, paw tremors, and grooming). Each behavior was defined and labeled frame-by-frame to align with the DeepLabCut data and to ensure standardization across LUPE recordings. BORIS provided the output of behavioral labels, timestamps, and event duration to serve as supervised training data (Friard & Gamba, 2016). On average, BORIS annotation required 2.5 hours per 30-minute video. Disagreements were resolved through consensus review. In total, approximately 90 hours were spent on manual annotation.

### B-SOiD: Unsupervised Classification

We initially explored the dataset using B-SOiD, an open-source, unsupervised behavior segmentation tool, for classifying withdrawal behaviors (Hsu & Yttri, 2021). B-SOiD clusters and labels similar behavioral motifs based on pose-derived features (Hsu & Yttri, 2021). There were, however, repeated miscommunications and errors during setup, as the Python environment required to run B-SOiD was outdated and incompatible with current versions of Anaconda and Python installed on our lab’s systems. As a result, B-SOiD was excluded from the final pipeline.

### A-SOiD: Classification

Pose data from DeepLabCut was turned into a binary table that captured how the mouse was moving in each video frame. This data was paired with BORIS-labeled behaviors which were used to teach the model which movement patterns corresponded to specific behaviors and to initialize training in A-SOiD. A-SOiD was then able to begin supervised learning on the known behavior classes. It is crucial to note that these behavior categories were defined by the experimenter during manual annotation, indicating the model was only able to learn what it had been explicitly shown. As these categories were predetermined based on human observation, this step introduced the potential for bias into the training process which is a limitation inherent in supervised learning approaches.

After supervised learning is complete, A-SOiD should employ its active learning strategy whereby the model identifies the frames it is uncertain about, typically those with low-confidence predictions, and flags them for human review. These frames should then be relabeled or confirmed by the experimenter, and the corrected labels should be added back into the training set for model refinement. In our study, the model completed 60 supervised learning iterations to identify and flag samples that it was uncertain about. However, due to technical issues in the current build of the software, the code failed to trigger the active learning correction step, preventing the model from identifying and refining samples it was uncertain about. This limitation restricted the classifier’s ability to improve over time and limited performance, particularly for ambiguous or rare behaviors.

It is important to note, that the model still classified ambiguous behaviors under an “other” category, however, the same technical issue prevented the software from running unsupervised learning on these behaviors. As a result, behaviors which were pre-defined that it was uncertain about may have been clumped into the “other” category, without an opportunity for the expert to manually refine familiar and discover novel behavioral motifs. While the active learning module holds potential for future studies, it is not yet a complete substitute for expert-guided classification.

## Results

We began by examining the temporal and frequency characteristics of opioid withdrawal behaviors in mice. The majority of withdrawal behaviors were brief, occurring between 1-2 seconds (**Fig. 5B**). Natural behaviors such as walking, being still, and rearing were the most common, while genital licking and grooming were the most frequently observed withdrawal-related behaviors (**Fig. 5B**). In contrast, jumping and paw tremors were less frequent and noticeably shorter in duration (**Fig 5B**). As a result, the distribution of behaviors was significantly skewed, introducing a class imbalance that made it difficult for the machine learning model to train accurately (**Fig 5A**) (Ghosh et al., 2024).

**Figure 5.**
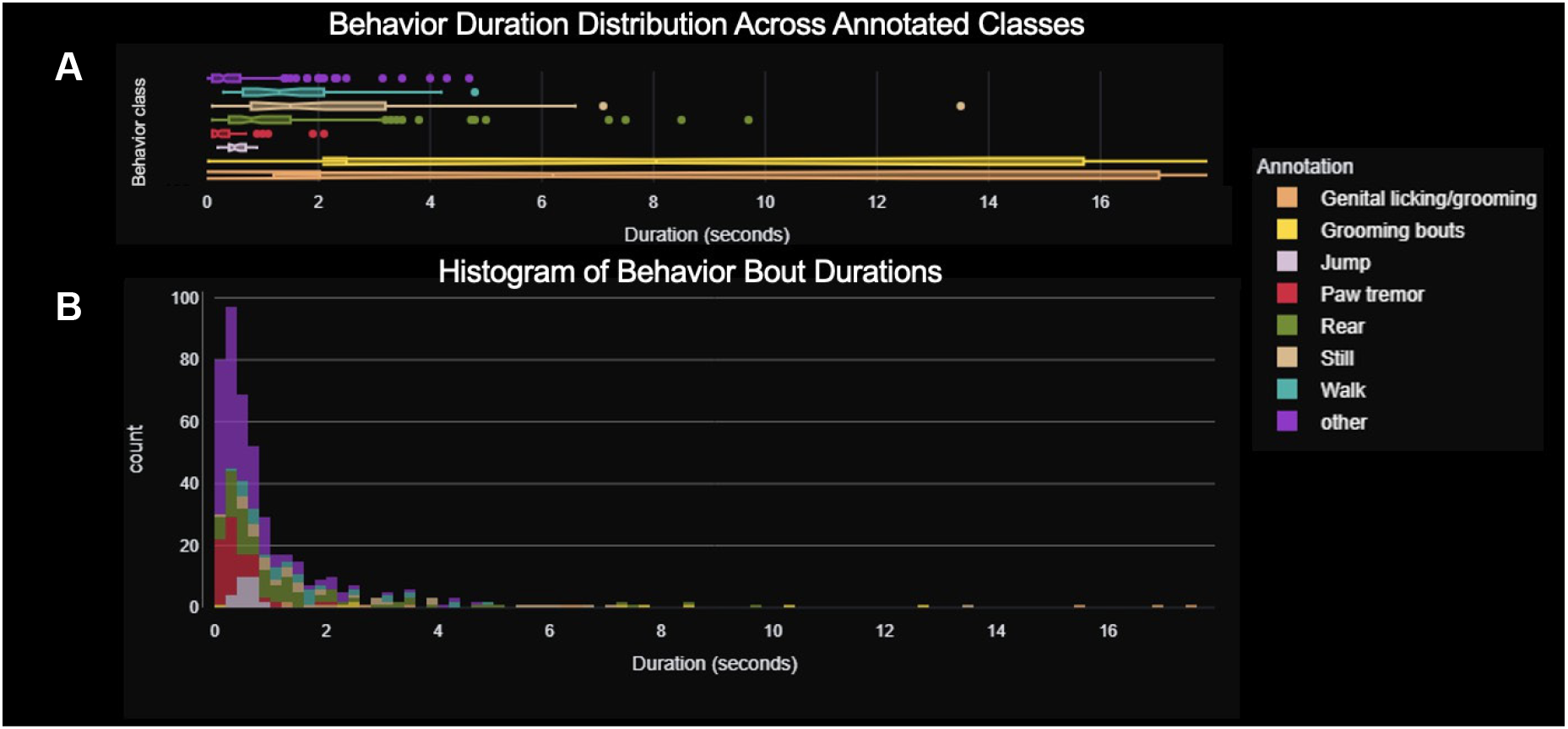
Behavioral Frequency and Duration. Duration distribution for each annotated withdrawal behavior, with grooming and genital licking bouts lasting the longest (**5A**). Behavior bout durations highlighting that most behaviors occurred in brief episodes under 2 seconds (**5B**).

This pronounced class imbalance presented a major challenge for behavior classification. To address this imbalance, we manually identified and annotated additional examples of underrepresented behaviors, such as jumping and paw tremors, by incorporating more videos that captured these actions. Originally, each behavior was portrayed with 10 samples, so we incorporated 30 examples of jumping and paw tremors to provide more data for the classifier to learn from. However, these additions did not substantially improve classification performance. This was not due to the number of examples available, but rather to the extremely brief duration of these behaviors. Jumping and paw tremors often occurred over just a few frames, far below the ∼400 frames per second at which A-SOiD performs best (Tillmann et al., 2024). As a result, the model had insufficient temporal information to learn robust features for classification. In contrast, behaviors such as genital licking and grooming typically lasted over 10 seconds, providing the model with more stable and informative data from which to learn (**Fig. 5A**).

Compounding this issue was the fact that A-SOiD’s active learning module could not be utilized due to technical issues. An error in the code prevented the software from advancing beyond the initial supervised training stage, blocking its ability to prompt review of uncertain frames. As a result, the classifier was unable to iteratively improve its performance after supervised training, failing to implement a core strength of the A-SOiD platform in this study.

Despite these constraints, the supervised model achieved moderate classification accuracy for high-frequency behaviors. The confidence of the model had a steady improvement across 60 iterations of active learning, as indicated by the increasing performance curves (**Fig. 6**). Behaviors that occurred with high frequency, including grooming, genital licking, walking, being still, and rearing, reached between 75-85% accuracy (**Fig. 6**). On the other hand, low-frequency behaviors, such as jumping and paw tremors, had higher variability between iterations as well (**Fig. 6**). Behavior-specific confidence scores reflect this discrepancy, with rare and short behaviors consistently receiving lower classification confidence. This suggests that the model relied on both sample size as well as duration per behavior for confident classification.

**Figure 6.**
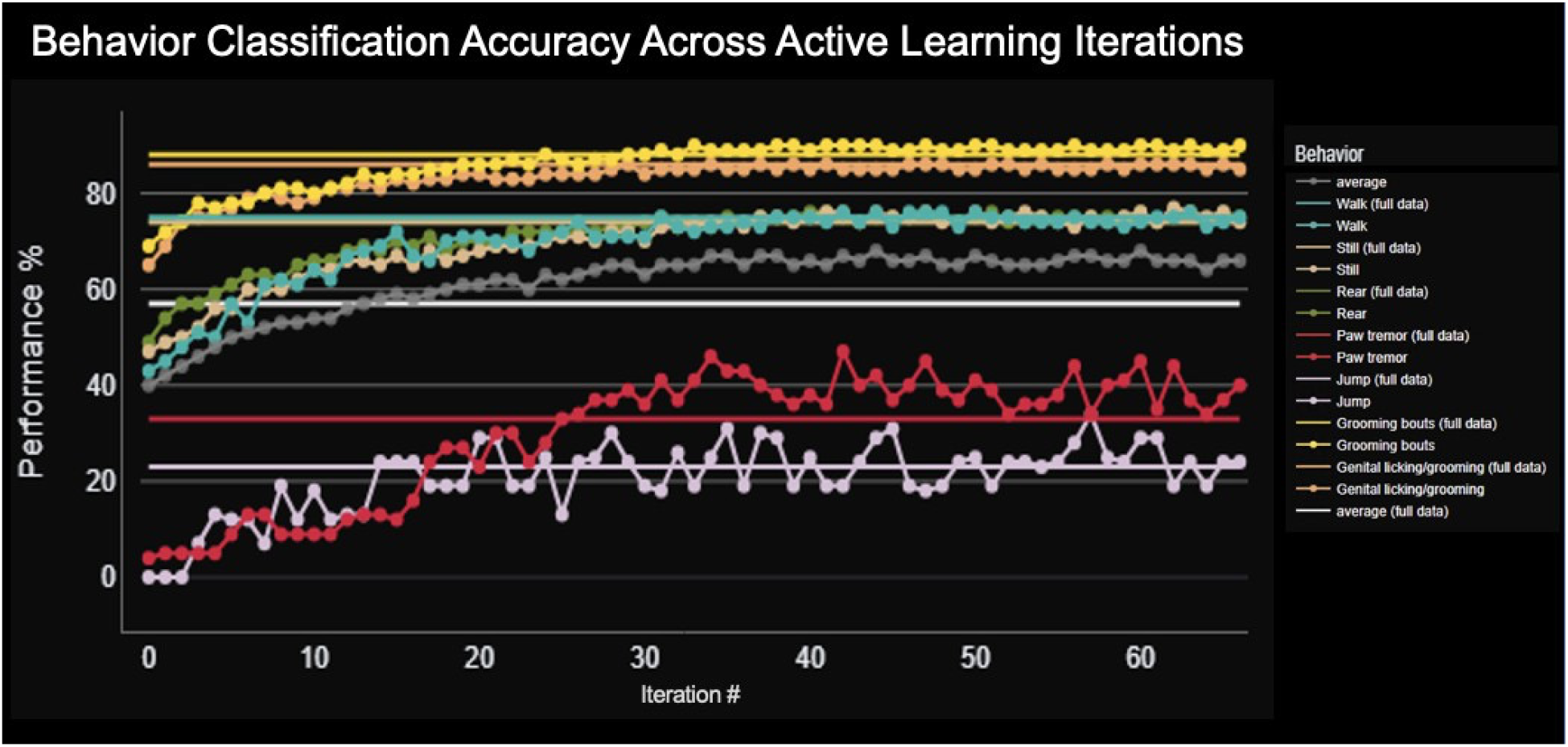
Model Accuracy Across Active Learning Iterations. Line plot showing classification accuracy of the A-SOiD behavioral model across 60 active learning iterations. Accuracy increased with each round of training, with performance improvements plateauing around iteration 40, indicating convergence of the model on an optimal feature space for classification.

The accumulation of training samples across model iterations revealed similar patterns: annotations for common behaviors like grooming and rearing increased rapidly, while annotations for jumping and paw tremors grew at a slower rate (**Fig. 7**). However, it is important to note that the total number of jumping and paw tremor events was not inherently limited as these behaviors were present in the data but were difficult for the model to identify confidently due to their brevity. For long-duration behaviors like grooming and genital licking, the model accumulated hundreds of examples, enabling robust learning. In contrast, the significant reduction in sampling for rapid behaviors suggests that these were either misclassified, absorbed into the “other” category (**Fig. 5**), or too short of a duration for the model to isolate and learn from. This pattern reflects a core limitation of applying standard deep learning pipelines to fast-onset, subtle, and short-duration behaviors,

**Figure 7.**
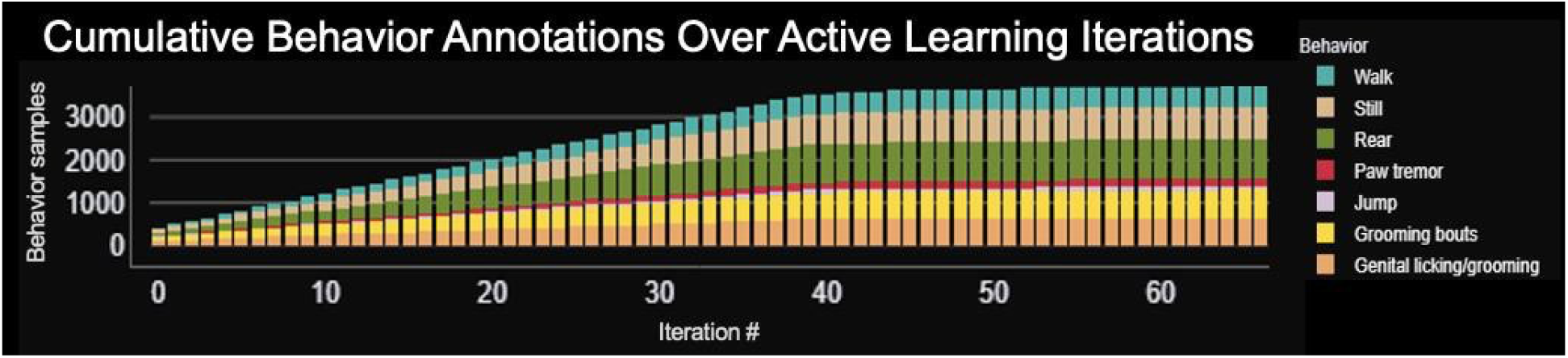
Cumulative Behavior Annotations for Active Learning. Total number of behavior annotations accumulated over 60 A-SOiD iterations. Grooming and genital licking behaviors dominated the labeled dataset, highlighting both their frequency and salience during opioid withdrawal in the annotated video segments.

Once supervised learning was completed, the software was unable to progress to the active learning stage due to an error in the code that prevented the A-SOiD interface from identifying uncertain behaviors and video samples for expert review. Consequently, the model could not iteratively improve its classification accuracy without the refinement loop nor could it implement its active learning strategy.

Critically, this same error prevented the interface from progressing to unsupervised learning to identify and label novel withdrawal behaviors in the “other” category (**Fig. 5**). As a result, the “other” category may serve as a source for noise and uncertain behaviors, rather than providing insight into mouse opioid withdrawal behaviors that may be missed.

## Discussion

This study demonstrates the feasibility and utility of using automated machine learning tools to classify behavior in clinically relevant animal models of opioid withdrawal, offering pronounced improvements in reproducibility and scalability over traditional manual scoring approaches. By integrating LUPE chambers for standardization of videos, BORIS for manual annotation, DeepLabCut for pose estimation, and A-SOiD for behavioral classification, we developed a pipeline that can be used to identify a subset of opioid withdrawal behaviors in mouse models. However, this process revealed several technical, conceptual, and methodological limitations for using machine learning to automate behavioral analysis, highlighting the importance of human expertise in future behavioral neuroscience studies.

Initially, we attempted to use B-SOiD, an unsupervised tool, to cluster similar behaviors based solely on motion features without requiring human annotations. Although this software has proved to be valuable in other studies for discovering and clustering behavioral motifs, we were unsuccessful in implementing it in our project. Despite multiple attempts over the course of the semester, we were unable to initiate behavioral clustering processes given outdated dependencies and a lack of recent maintenance for the software. This highlights a common issue in the field: open-source tools often become unusable if they are not actively supported, despite their initial promise. It is vital to ensure that other tools, including A-SOiD, are supplied with long-term maintenance plans to ensure that they can be used across labs and over many years.

Nevertheless, the automation of behavioral scoring has important implications for accelerating research in a variety of fields. This advancement can significantly reduce the time required for manual scoring, which is labor-intensive and can cost labs thousands of dollars to complete. Not only does automated behavioral classification greatly reduce the resources needed per experiment, but it also allows for more precise and temporally aligned studies of brain-behavior relationships. In particular, many withdrawal behaviors in mice are believed to be driven by distinct circuits in the brain. For example, jumping is believed to be a high-energy, escape response to a stressful stimulus while paw tremors may be the result of a dysregulation in the autonomic system of a mouse—these distinct behaviors highlight the possibility that different brain regions are involved in an array of withdrawal symptoms (Siegel et al., 1975; Kim & Im, 2024). Therefore, matching precise timepoints with neural activity recorded through a variety of techniques, including calcium imaging, fiber photometry, or electrophysiology, will allow researchers the ability to begin to unravel the neural circuitry involved in specific withdrawal behaviors. This, in turn, could propel research forward in developing treatments that target specific symptoms of withdrawal from opioids.

Our current model shows moderate success for classifying natural and withdrawal-related behaviors that last for relatively long durations of time (**Fig. 5**). However, the classifier showed difficulty in identifying brief events, suggesting that current open-source platforms may not be equipped to capture and analyze rapid and nuanced behaviors like paw tremors. As a result, we could not initiate active learning on this dataset. It is also important to note that our model ran on supervised learning, rather than active learning, which creates an inherent human bias present in the data analysis. Although our study did not reach its full potential due to software limitations, we developed the infrastructure for future work to be conducted in this field. With updated open-source tools, improved processing for shorter frame rates, and functional active learning strategies, this pipeline has the potential to serve as a reproducible platform for modeling a wide range of affective behaviors, including withdrawal and pain, across conditions and species. It is essential for collaboration to occur in the development of these tools, allowing for enhanced cross-lab validation and the generalization of open data standards to propel the field of behavioral neuroscience forward.

### Future Directions

To address the challenges uncovered in our project, we plan to implement several refinements to our behavioral analysis pipeline. In particular, to overcome the difficulty in classifying behaviors of brief duration, we plan to increase the temporal resolution of our recordings with cameras that capture higher frame rates. This will allow us to accurately track fast and subtle behaviors that may be underrepresented in datasets by ensuring that rapid movements are visible across more frames. In turn, this will provide A-SOiD with more information to train on and learn from. Additionally, future versions of our pipeline will make use of A-SOiD’s active learning module after the current interface errors are resolved. This will allow us to accurately flag uncertain video segments for refinement and correction. Consequently, active iterative refinement will reduce dependence on initial annotation quality and quantity, making behavioral analysis more efficient and robust and reducing the need for intensive manual scoring. Furthermore, we intend to implement unsupervised learning to explore potentially novel behavioral motifs captured in the “other” category, with the potential to determine the variety of withdrawal behaviors that animal models can exhibit after opioid withdrawal is precipitated.

These improvements are necessary to proceed with integrating behavior with neural activity. By aligning timestamps of behavioral events with calcium imaging, fiber photometry, or optogenetic manipulation, we may be able to disentangle the relationship between specific withdrawal behaviors, such as jumping or genital licking, and neural data. This will allow us to identify which brain regions and circuits are involved in specific features of withdrawal, enabling researchers to develop precise interventions for these behaviors.

In addition to this, we plan to expand the dataset to include a larger sample of animals, more annotators, and different experimental conditions. Adapting this model for the administration of oxycodone or using rats as models will extend the pipeline’s utility across domains of behavioral neuroscience. Overall, we plan to develop an open-source reproducible behavioral classification network that can be validated and implemented across labs and for an array of conditions in the years ahead.

## Conclusions

Currently, our study presents a high-resolution, scalable, and partially automated behavioral pipeline for classifying and quantifying withdrawal behaviors in mice through machine learning. Through LUPE video acquisition, DeepLabCut pose estimation data, BORIS manual annotations, and A-SOiD machine learning classification, we developed a framework which is capable of identifying withdrawal and natural behaviors in mouse models with moderate success. The pipeline demonstrated relatively robust performance in identifying long-duration behaviors, like genital licking, while uncovering the limitations of current classification methods in detecting rapid events, like paw tremors.

Although there were technical setbacks, particularly in the inability to initiate the active learning and unsupervised clustering functions in A-SOiD, our study lays the groundwork for the use of machine learning in future behavioral analysis. This project suggests that machine learning tools can reduce human labor, improve reproducibility, and enhance data analysis while also indicating that human expertise and long-term maintenance of open-source platforms are critical in advancing the field of behavioral neuroscience.

Importantly, our study has set the stage for future studies that aim to link behavioral events with neural activity. We intend to apply this pipeline to an array of behaviors and experiments, such as those studying pain-related behaviors or drug-seeking activities across species and conditions. This project exemplifies the promise and challenges of using machine learning to model and analyze behavioral states, however, with collaborative refinement and improved tools, we anticipate that there will be more precise, scalable, and advanced neuroscience research in the years to come.

## Acknowledgements

I would like to thank Dr. Gregory Corder and the entire Corder Lab for their mentorship and support. Special thanks to Lisa Wooldridge and Justin James for their guidance and feedback as well as Alex Hsu and Dr. Eric Yttri for their development of the A-SOiD platform. This project was supported by the National Institute of Health and the National Institute on Drug Abuse as well as the 5R01DA056599-03 grant for ‘Circuitry dynamics underlying opioid-dependence: Integrating structural, functional, and transcriptomic mechanisms.’

## Author Contributions

Conceptualization: SAC, LMW, GC

Methodology: SAC, LMW, JJ

Formal analysis: SAC

Investigation: SAC, LMW, JWKW

Writing: SAC, LMW, GC

Supervision: LMW, GC

